# Extended *t*-cores for the *de novo* identification of transposable elements and other inexact repeats from short read RNA-seq data

**DOI:** 10.64898/2026.07.06.736737

**Authors:** Sasha Darmon, Arnaud Mary, Vincent Lacroix

**Author notes:** Equal contribution.

## Abstract

Transcribed repeats represent a major challenge in the *de novo* assembly of transcriptomes from short RNA-seq reads. Young transposable elements (TEs) and more generally, inexact repeats, create dense and ambiguous regions in the assembly graph, preventing the correct assembly of transcripts. In this paper, we introduce a fully *de novo* method based on the discovery of dense regions in the compacted De Bruijn graph (DBG) to identify such repeats directly from short read RNA-seq data, without requiring a reference genome or repeat database. Our approach defines the *extended t-cores*, subgraphs of the DBG that capture the complex topology induced by expressed inexact repeats appearing in RNA-seq reads.

Independently of its interest for transcriptome assembly, the proposed method appears to be effective for the *de novo* identification of repeats in transcriptomes. After classifying cores using sequence-based motifs to distinguish simple repeats from potential TEs, we demonstrate its potential for the *de novo* discovery of TEs.

We validate the approach on a *Mus musculus* dataset, showing that extended *t*-cores correspond to known expressed TE families. We also illustrate its *de novo* discovery potential on a non-model species, *Canis lupus familiaris*, where the method was also able to recover known TEs.

## Introduction

In 2026, the NCBI Sequence Read Archive (SRA) (Leinonen et al., 2010) database has accumulated more than twelve million Illumina short-read RNA-seq datasets, collected from tens of thousands of unique species and a wide variety of tissues. Even though these datasets are freely available, their full potential remains clearly under-explored.

The analysis of such data follows two main paradigms (Haas and Zody, 2010). Either RNA-seq reads are mapped onto a reference genome, or they are assembled *de novo*. Most projects favour the mapping approach taking advantage of the growing availability of reference genomes. Despite recent efforts to standardise genome assembly (Lawniczak et al., 2022), the quality of these reference genomes is however still uneven and many genomes are only assembled at the contig or scaffold level, leaving repeated regions poorly assembled, including their associated polymorphism. RNA-seq reads stemming from these repeated regions are therefore undetected, which leads to an underestimation of the prevalence of repeats in transcriptomes.

Transcriptome assembly, on the other hand, assembles the reads independently of the genome (Hölzer and Marz, 2019). All reads participate in the assembly, including the ones which stem from repeats. These reads create ambiguities in the assembly process. When a repeat corresponds to a subsequence of several genes, the assembler can produce two types of errors: either it stops assembling and produces truncated transcripts, or it continues to assemble, but following the wrong path and produces chimeric transcripts.

*De novo* assembly of full-length transcripts is therefore particularly challenging when transcripts span repetitive regions. Focusing on the simpler objective of identifying the repeats themselves seems more tractable and would already constitute valuable information for transcriptome assembly. It is also highly relevant to the community of researchers interested in repeats.

Among repeats, transposable elements (TEs) are of particular interest and are quite problematic to study. They correspond to sequences that can copy and insert themselves throughout a genome. In maize, TE copies correspond to up to 85% (Baucom et al., 2009; Schnable et al., 2009) of the genome. The vast majority of these TE copies have lost the ability to transpose, as they have accumulated mutations thoughout evolution. Out of all copies, studying the ones that are expressed is of particular interest, because they are more likely to have an impact on the phenotype. Some of the expressed copies may still be able to transpose and therefore generate new insertions. Those which have lost the ability to transpose may still have an active promoter and therefore impact the expression of genes located nearby possibly generating gene-TE chimeras ((Lanciano and Cristofari, 2020)). Finally, those which do not have an active promoter may still be transcribed, if they are hosted within transcribed genes. Out of those hosted within transcribed genes, very few insertions are located in coding exons, some are in UTRs and many are in introns. RNA-seq data corresponds in vast majority to processed RNA sequences where all introns have been fully spliced. However, even when using stringent protocols which focus on poly(A)+ transcripts, there always remains a fraction of ∼ 5% of pre-mRNA (Tilgner et al., 2012) which explains why intronic TEs can also generate RNA-seq reads.

Importantly, our ability to discriminate two copies of a TE depends on the "age" of the TE. While old copies are easy to distinguish thanks to the accumulation of mutations, newly transposed copies are identical to their original template, and therefore impossible to distinguish at the read level. The types of repeats that are most problematic for transcriptome assembly are high-copy-number low-divergence repeats.

Analysing the expression of TEs with short reads requires dedicated bioinformatics methods (Lanciano and Cristofari, 2020). First, using repeat databases (such as Dfam (Hubley et al., 2016) or RepBase (Jurka et al., 2005)) enables to obtain consensus sequences for TE families, mapping RNA-seq reads to these consensus (for instance, using *TETools* (Lerat et al., 2017)) enables to estimate the expression level of the TE family. Second, using reference-based genome annotations, it is possible to try to estimate the contribution to the expression of each genomic copy using tools such as TETranscripts (Jin et al., 2015) or SQUIRE (Yang et al., 2019). Yet, this second strategy is often unfeasible for non-model species where well annotated genome assemblies are not always available. Third, applying *de novo* genomic repeat annotation tools (such as Repeat Explorer (Novák et al., 2013), REPdenovo (Chu et al., 2016), Tedna (Zytnicki et al., 2014), RepARK (Koch et al., 2014), dnaPipeTE (Goubert et al., 2015)) to RNA-seq data; however, these approaches have been designed for genomic data, and applying them to transcriptomic data carries a high risk of bias due to the variability of transcript expression levels.

In this paper, we address the general problem of *de novo* identification of inexact repeats from short RNA-seq reads. Inexact repeats refer to highly similar but divergent sequences within the transcriptome, which include transposable elements (TEs) and simple repeats. Our method operates fully *de novo*, without a reference genome, or a repeat database. It can therefore be applied to any species. Our method is simple as it only uses the structure of the de Bruijn graph. Yet, it gives promising results, opening avenues for further developments.

Since nodes of a De Bruijn graph have a degree bounded by the size of the alphabet (here 4), we define the "extended degree" of a *k*-mer as the number of distinct *k*-mers reachable at a fixed distance, in order to represent the number of locally distinct transcripts containing that *k*-mer. We then compute the extended *t*-cores of the graph as the maximal connected subgraphs containing only nodes of extended degree higher than *t*. Specifically, this method targets transcribed inexact repeats, the very elements that create dense regions in the assembly graph and hinder transcript reconstruction. By design, this topological approach will not recover low-copy or perfectly identical TEs, as they lack the structural complexity to form such cores. Finally, we propose a classification of the identified repeats and show the potential of our method for the *de novo* discovery of TEs.

Our implementation demonstrates linear time and space scalability, as each step, including the BFS-based graph traversal, operates in linear time with respect to the number of nodes in the compacted De Bruijn graph. Furthermore, we leverage BCALM2 (Chikhi et al., 2016) to construct the compacted De Bruijn graph with high efficiency and low memory overhead. As a performance benchmark, our tool processes a dataset of one hundred million reads in less than an hour on a standard laptop. The software is available on GitHub: https://github.com/sdarmon/ET-core.

To evaluate our method, we first benchmarked it on a *Mus musculus* dataset, where TEs are well characterised and consensus sequences are available in Dfam. We find that 92% of our predictions indeed match TE consensus. Conversely, 17% of TE consensus identified as expressed by TETools are found by our method. We then turn to the analysis of a *Canis lupus familiaris* dataset, where Dfam does not contain any TE consensus, in order to demonstrate its potential for the *de novo* identification of TEs. Again, we find that more than 90% of the extended-t-cores we predict as potential TEs do correspond to TEs. Importantly, we make these predictions without a reference genome or repeatome.

Finally, we would like to insist on the fact that the method we propose enables to analyse repeats in transcriptomes using very limited resources. It runs in less than an hour on a desktop computer, even with hundreds of millions of reads. More importantly, it can be run using existing short reads RNA-seq data and does not require to produce new long read data. Restraining from producing new data is a true challenge since the carbon footprint of research in most labs is dominated by purchases (De Paepe et al., 2024).

## Material and methods

### Datasets

We used two RNA-seq short-read datasets: one from ***Mus musculus*** (Sessegolo et al., 2019) (brain cells, model species, used for method evaluation) and another from ***Canis lupus familiaris*** (Prouteau et al., 2022) (oral skin cells, non-model species, used for the *de novo* identification of TEs). In both datasets, only poly(A)+ RNAs were sequenced.

The reads were preprocessed using *FastP* (Chen et al., 2018) to remove adapters, poly-A/T tails, and low-quality sequences. This common step is crucial to ensure that these sequences are not misidentified as repeated content in the transcriptome.

Furthermore, homopolymers were compressed to a maximum of five nucleotides. This strategy, frequently used to improve mapping and assembly performance ((Blassel et al., 2022), e.g. in the aligner *Minimap2* (Li, 2018), and in the assembler mDBG (Ekim et al., 2021)), allows the analysis of repeats that span across long homopolymeric runs.

### Computational Resources

All benchmarks and computations were performed on a personal laptop running Ubuntu 24.04.4 LTS. The hardware consisted of an Intel® Core™ i5-1335U processor (10 cores: 2 Performance at up to 4.6 GHz, 8 Efficient at up to 3.4 GHz; 12 threads), 16 GB of single-channel DDR5-5200 MHz RAM, and a 512 GB Western Digital PC SN740 PCIe 4.0 NVMe SSD. To ensure consistent performance, the system was connected to AC power with the OS CPU power governor set to ’performance’.

The source code is written in *Rust*, *Python 3* and *C++* and is available on GitHub : https://github.com/sdarmon/ET-core through a single bash script. Refer to the README for more detailed explanations for used programs.

## Methods

In this section, we explain how we compute extended *t*-cores and how we derive our *de novo* repeat classification.

### Assembly graph: compacted De Bruijn graph

Given an integer *k*, a De Bruijn graph (DBG) represents every *k* − 1 overlap between the *k*-mers, i.e. the reads’ substrings of length *k*. More precisely, an edge connects two *k*-mers if they share an exact overlap of *k* − 1 nucleotides. We specifically chose to work with the compacted version of the De Bruijn graph, cDBG, where every non-branching path is represented as a single node with a corresponding compacted sequence called a unitig. While having the same structure as the DBG, the cDBG has multiple benefits: it can be efficiently computed from the *k*-mers of RNA-seq reads, it is highly space-efficient, and it is widely used for sequence assembly (Chikhi et al., 2016; Compeau et al., 2011; Huang et al., 2023).

More importantly for our approach, (c)DBG naturally collapses repetitive sequences into highly connected, dense regions, provided they share at least one common *k*-mer. While these dense regions traditionally represent a major bottleneck for sequence assembly algorithms, our extended *t*-core approach specifically exploits these graph structures to directly identify inexact repeats.

Yet, a major challenge when working with De Bruijn graphs is that sequencing errors create false branching and spurious structures within the graph. Since we do not want to capture these irrelevant structures, we filter out sequencing errors by removing nodes in the graph that have an abundance smaller or equal to a threshold *a*. In practice, since we work with transcriptomic data, the abundance oaf valid k-mers can be low for poorly expressed genes, hence we choose *a* = 2. Conversely, sequencing errors located in highly expressed genes may be supported by many k-mers. We therefore also remove nodes supported by an abundance which is 20 times lower than its predecessor. This relative coverage threshold can be considered as conservative since the sequencing error rate for Illumina platforms is typically less than 1% (Pfeiffer et al., 2018; Stoler and Nekrutenko, 2021).

### Definition of the extended degree of a unitig

We define the extended degree of a unitig in the cDBG as the number of distinct *k*-mers reachable within a fixed distance of *n* = 10 nucleotides. The value of *n* was chosen to allow a sufficient number of divergent paths (up to 4^10^ ≈ 10^6^) to represent the distinct transcripts containing that unitig.

Simultaneously, keeping *n* relatively small compared to *k* allows a thin granularity, avoiding cores encompassing all adjacent repeats. This double constraint ensures that our method captures the inexact repeats transcribed within different genes, which constitute the main bottleneck in *de novo* assembly.

Despite the precautions taken against sequencing errors, some of those can still artificially increase the extended degrees. That is specifically the case for the sequencing errors within homopolymers, where the reachable *k*-mers differ by only one or two substitutions. Indeed, since homopolymers are compressed to a maximum of five nucleotides during preprocessing, any sequencing error *Y* occurring within a long homopolymer of nucleotide *X* (length *L*) will generate an identical compressed sequence (e.g., *X* ^5^*YX* ^5^), creating a false branch in the graph. Because this error can occur at multiple positions along the original homopolymer, the abundance of such erroneous *k*-mers multiplies by up to a factor *L* − 10, risking to exceed our relative coverage threshold. To systematically eliminate the influence of these sequence errors, we refine our definition: the extended degree of a unitig is defined as the number of equivalence classes of distinct *k*-mers reachable within *n* nucleotides. These classes are formed by the transitive closure of the Hamming distance relation, where two *k*-mers are grouped together if their Hamming distance is no greater than 2. In practical terms, this step acts as counting the number of connected components of a graph, where the nodes are the *k*-mers, and where two *k*-mers are connected if they have ≤ 2 mismatches. This allows much more divergent *k*-mers to be grouped together, provided there is a continuum of low divergent *k*-mers linking them.

### Computation of the threshold t

To differentiate unitigs with a high extended degree from those with a low extended degree, we must define a specific threshold, *t*. We consider two modes.

In the SENSITIVE mode, the threshold *t* is defined as the lowest integer such that the high extended degree unitigs (i.e., those with a degree ≥ *t*) represent less than 1% of all unitigs in the graph. In the PRECISE mode, *t* is defined as the lowest integer such that high extended degree unitigs represent less than 0.1% of all unitigs.

The purpose of the PRECISE mode is to strictly limit the number of high extended degree unitigs, thereby maximising the precision of the captured inexact repeats and reducing false positives. Conversely, the SENSITIVE mode applies a more permissive threshold, aiming to maximise the recall (sensitivity) of inexact repeats, albeit at the cost of a slightly lower precision.

### Extended t-core computation

Once the threshold *t* is established, the algorithm consists of a breadth-first search (BFS) starting from the highest extended degree nodes to extract the extended *t*-cores from the compacted De Bruijn graph. Formally, an extended *t*-core is defined as a maximal connected component within the subgraph induced exclusively by high extended degree unitigs (i.e., unitigs with an extended degree ≥ *t*).

Depending on the size of the repeat, the extended *t*-cores either capture the full sequence (for short repeats such as microsatellites) or only the boundaries of the repeat (for longer elements such as TEs). This structural difference is explained by the uneven distribution of the extended degree along long repetitive sequences. Unitigs located at the boundaries of a repeat exhibit a high extended degree because their reachable *k*-mers (at a distance of *n* = 10 nucleotides) diverge into the various distinct host genes or flanking genomic regions that contain the repeat. Conversely, unitigs located in the middle of a long repeat exhibit a low extended degree: their reachable *k*-mers at distance *n* remain strictly internal to the repeat consensus, giving highly similar *k*-mers. Moreover, a representative sequence is extracted for each extended *t*-core. For a given core, we define the representative sequence as the most abundant path through its unitigs (i.e., the path with the highest cumulative read coverage). This sequence can serve as a *de novo* consensus for that core providing a template for downstream analysis.

### De novo repeats classification and validation

To facilitate the biological interpretation of the identified repeats, we implemented a *de novo* classification that characterises each extended *t*-core based on its sequence content and its structural context within the global assembly graph.

First, we define two simple repeat motifs that can easily be identified only with their sequences :

- A microsatellite is defined as a motif of 2 to 20 nucleotides, repeated at least twice, and spanning a minimum of 8 consecutive nucleotides.
- An X-stretch is defined as a single nucleotide consecutively repeated at least five times, typically corresponding to collapsed homopolymers.

Then, we use those definitions to divide the extended *t*-cores into two categories depending on the basic repeat motifs identified: simple repeat or Potential TE.

The simple repeats category comprises: *(i)* X-stretch-rich cores (defined as cores where the total number of identified X-stretches is ≥ 0.75*u*, with *u* being the number of unitigs within that core), and *(ii)* microsatellite-dominated cores (defined as cores where microsatellite motifs account for at least 50% over all sequences, or where at least 50% of the unitigs individually consist of ≥ 50% microsatellite motifs).

Conversely, the Potential TEs category contains all remaining extended *t*-cores that do not fall into the *simple repeats* classification. We focus on Potential TEs extended *t*-cores to evaluate the method for identifying TEs.

To validate the method, we used the *Dfam* repeat database (Hubley et al., 2016) to retrieve consensus sequences of repeats specific to the lineage of the studied species. For each species, we collected from *Dfam* the curated consensus sequences classified as Interspersed Repeats; TEs that were assigned to the species, or its ancestors. Next, we quantify the number of expressed TEs, by following *TETools* (Lerat et al., 2017) pipeline. Reads were aligned to the repeat consensus sequences using *Bowtie2* (Langmead and Salzberg, 2012) with the default option, and reads were counted using *FeatureCounts* (Liao et al., 2014). Reads with less than 50% of their length overlapping a consensus were discarded. We consider a TE consensus to be expressed if it exhibits a mean coverage depth greater than 1 and a breadth of coverage greater than 50%.

Then, we define true positive cores as those containing at least one unitig that successfully aligns to an expressed TE consensus (according to the previous definition). To do so, we aligned all unitigs using Bowtie2 with default scoring parameters, which typically tolerate less than a 10% mismatch rate, ensuring high alignment confidence. However, to account for the natural evolutionary divergence between transcribed TE copies and their reference consensus sequences, we decreased the seed length parameter (and adjusted the seeding sensitivity). This modification doesn’t affect the alignment score. It only allows to recover valid alignments that otherwise would not have been tested at all, due to localized mismatches in the seeds. Indeed a single mismatch in a short unitig (41 nt) can dismiss all potential seeds (22nt).

## Results

### Extended *t*-cores and inexact repeats

The first extended *t*-core of the *Mus musculus* dataset (which possesses the highest extended degree) and its neighbourhood are depicted in Figure 1. While this core is composed of only 73 unitigs (nine of which are shown in Figure 2), it is connected to more than 170,000 other unitigs within a distance of less than 20 unitigs. The high extended degree of this core stems from its position at the 5’ end of the *B2_Mm1a* TE family, a young and active family with thousands of new and low divergent copies within the *Mus musculus* genome (Ichiyanagi et al., 2021).

**Figure 1.**
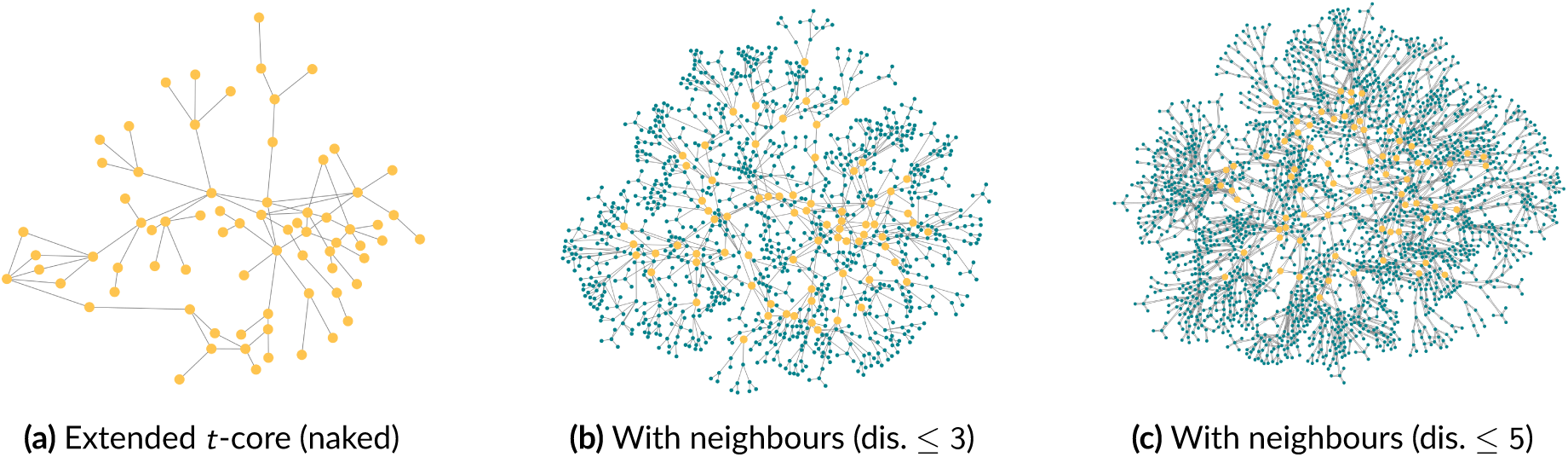
(a) First extended *t*-core (in yellow) of the *Mus musculus* dataset. (b-c) Its neighbors at distance 3 (b) and 5 (c) are depicted in blue. Nodes at distance 6 or more are not represented.

**Figure 2.**
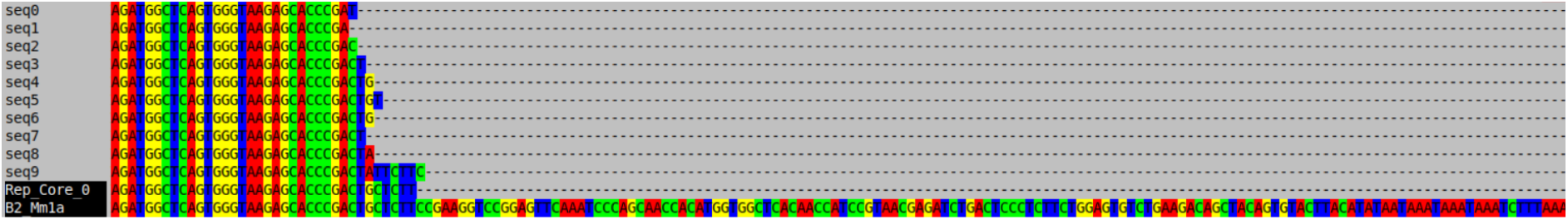
Alignment of ten unitigs from the first extended *t*-core and its representative sequence (Rep_Core_0) to the consensus sequence of the B2_Mm1a mouse TE.

Because these copies have not yet diverged significantly, many of them share k-mers, whereas divergent regions correspond to alternative k-mers, all linked to the shared one. Moreover, *SINE* families (such as B2) are distributed across thousands of distinct loci (Consortium et al., 2002; Ichiyanagi et al., 2021), which generate an exceptionally dense and complex subgraph within the assembly graph even if only a subset of those loci are transcribed.

This first core corresponds to the 5’end of the *B2_Mm1a* family. Beyond this first core, out of the 20 first cores, we find that 16 also match extremities of known families of TEs. Out of the 4 remaining, 3 correspond to simple repeats ((*AAAC*)*^n^*, (*AAAAC*)*^n^* and (*AAGG*)*^n^*), and 1 corresponds to a non coding RNA (*Snhg11*).

This suggests that our method could be used for the *de novo* prediction of TEs. In particular, cores which correspond to simple repeats can be flagged easily. Non-flagged cores are called Potential TEs (see section Methods).

Of note, out of the 20 first cores, core 15 corresponds to the 3’end of *B2_Mm1a*. Core 10 and 12 correspond to alternative versions of the 5’end of *B2_Mm1a,* possibly suggesting a sub-structure of the family (see Figure 3 and discussion).

**Figure 3.**
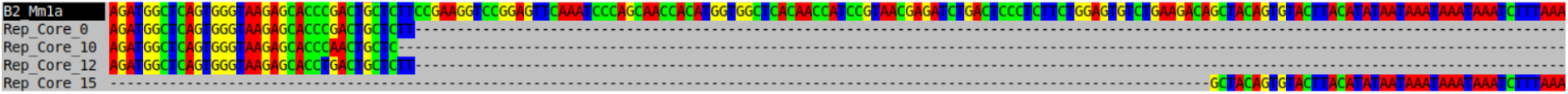
Alignment of the representative sequences of the extended *t*-cores 0, 10, 12 and 15 to the consensus sequence of the B2_Mm1a mouse TE.

In the next section, we evaluate the potential of our method to identify *de novo* TEs. We consider a core as a true positive when it matches a DFAM consensus.

### Extended *t*-cores for *de novo* identification of TEs

#### Performance on potential TE extended t-cores

Running the extended *t*-core algorithm in PRECISE mode (default mode, see Methods) on the *Mus musculus* dataset yielded 997 distinct cores, with the threshold *t* calculated to be 10. It is crucial to emphasize that these results were achieved entirely *de novo*, without any reliance on a reference genome or prior knowledge of TE sequences.

Out of the top 100 cores, 73 of them were assigned to "Potential TEs" and 71 (97.3%) were actual true positives (see Methods for exact definition). The two false positive cores correspond to *Snhg11*, a non coding RNA. Considering all the 997 distinct cores, 689 (69.1%) of them were assigned to Potential TEs and 637 were actual true positives leading to a precision of 92.4%. The recall for these 689 cores was 17.0%, meaning that among all true positives (TE consensus sequences considered as expressed using TETools), 17.0% were successfully captured within the "Potential TEs" extended *t*-cores in PRECISE mode.

It is noteworthy that the PRECISE mode provides high precision at the expense of sensitivity, a trade-off that is consistent with its design. Furthermore, when considering only the top-ranked extended *t*-cores (those with the highest maximal extended degrees), the precision is even higher, demonstrating that the most prominent signals captured by our approach consist almost entirely of true TEs, even though we did not use any TE references.

To better represent the precision and sensitivity trade-off, we plotted a cumulative performance curve by ordering the "Potential TEs" extended *t*-cores in descending order of their maximal extended degree. For each core at rank *i*, we define its cumulative precision as the proportion of the top *i* cores that match at least one TE consensus sequence. Similarly, the cumulative sensitivity (or recall) is defined as the ratio of unique TE consensus sequences recovered by the top *i* cores to the total number of expressed TE consensus sequences in the dataset.

The resulting performance curve is depicted in Figure 4. For the sake of completeness, we extended this analysis beyond the *t*-cores by including the remaining nodes of the graph, also ordered by their decreasing extended degree. The transition between the identified *t*-cores and the remaining graph nodes is indicated by a green vertical dashed line.

**Figure 4.**
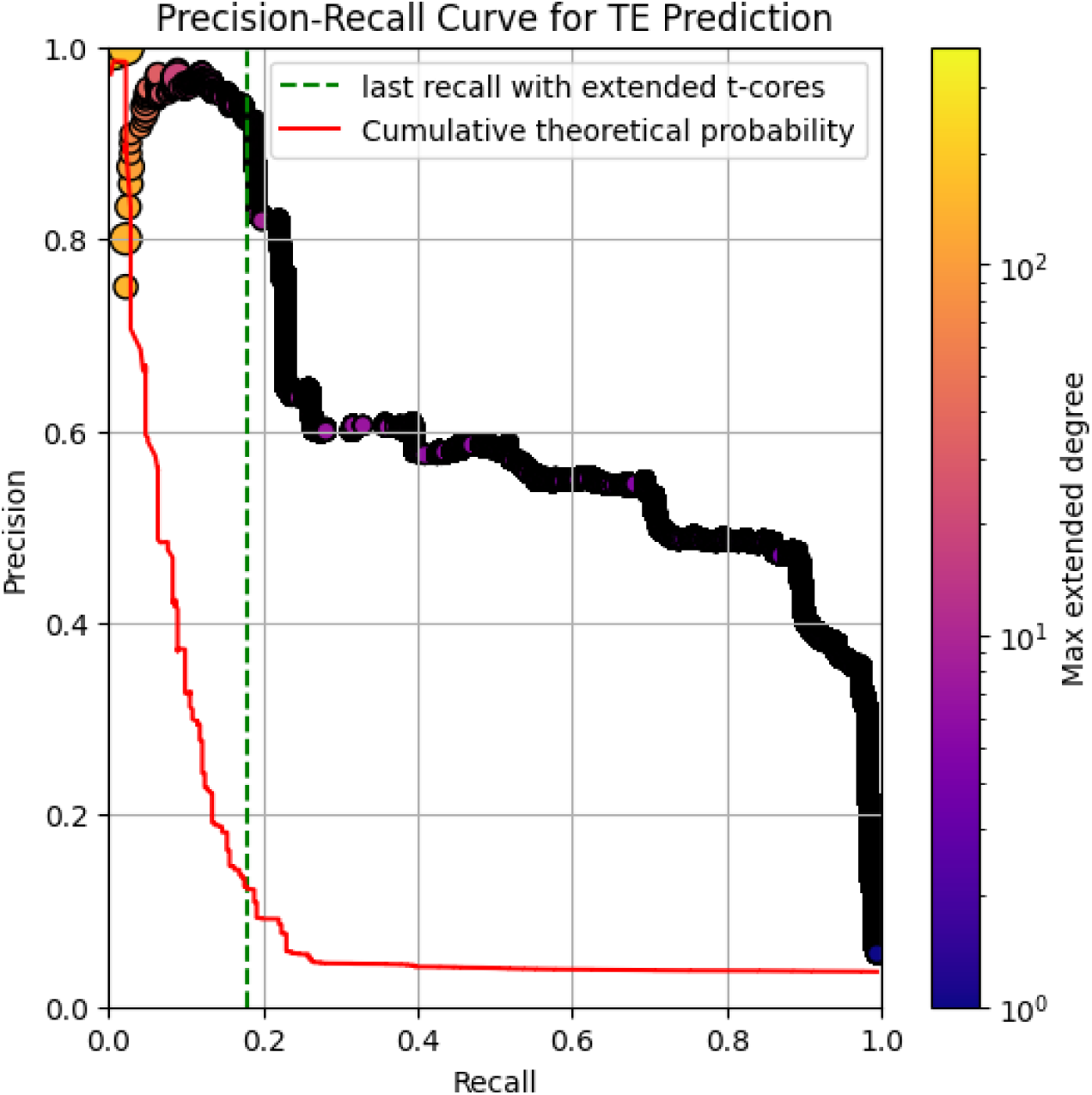
Cumulative performance restricted to "Potential TE" extended *t*-cores on the *Mus musculus* dataset. The graph shows the cumulative precision and sensitivity specifically for cores flagged as "Potential TEs", ordered by descending maximal extended degree. Dot sizes are proportional to the number of unitigs in each core. The red line represents the baseline performance expected from a random selection of unitigs, and the vertical green dashed line indicates the threshold *t* = 10 (PRECISE mode).

To assess the statistical significance of our results, we compared the cumulative performance of the extended *t*-cores against a random baseline. Since a single aligned unitig is sufficient to validate an entire core as a true positive, we generated a null model for comparison. This null model consists of randomly selected sets of unitigs from the graph, with the constraint that they have the same cardinality (size) as our identified *t*-cores (indicated by the red line in Figure 4).

As shown in Figure 4, this random selection strategy significantly underperforms compared to the extended *t*-core approach. The large gap between the two curves demonstrates the efficacy of our method in specifically isolating biologically relevant repeats rather than arbitrary graph structures, confirming that structural topology alone is a powerful predictor for TE discovery without any prior sequence knowledge.

Worth noting, the first drop of the cumulative precision at 75% is due to the fact that the fourth core is a false positive, corresponding to *Snhg11*, leading to a drop of 1/4th of the precision. The second drop, which brings the cumulative precision down to 60%, corresponds to the transition from evaluating full extended *t*-cores to analyzing individual unitigs with an extended degree of 13. At this threshold, a single unitig has a 60% probability of representing a TE; this probability steadily decreases as the extended degree becomes lower.

#### Performance of the SENSITIVE mode for TE identification

To evaluate the capacity of our method to perform a more exhaustive search of the TEs, we analyzed the *Mus musculus* dataset using the SENSITIVE mode (more permissive mode, see Methods). With the threshold established at *t* = 5, the algorithm identified a total of 13598 extended *t*-cores with 9987 (73.5%) cores classified as Potential TEs.

Over the first 1000 cores, 740 cores are classified as Potential TEs with a precision of 69.4% and a sensitivity of 33.2%. Considering all 9987 Potential TEs cores, the global sensitivity reaches 67.4%, while its precision is measured at 42.7%. A ROC curve is depicted on Figure 5.

**Figure 5.**
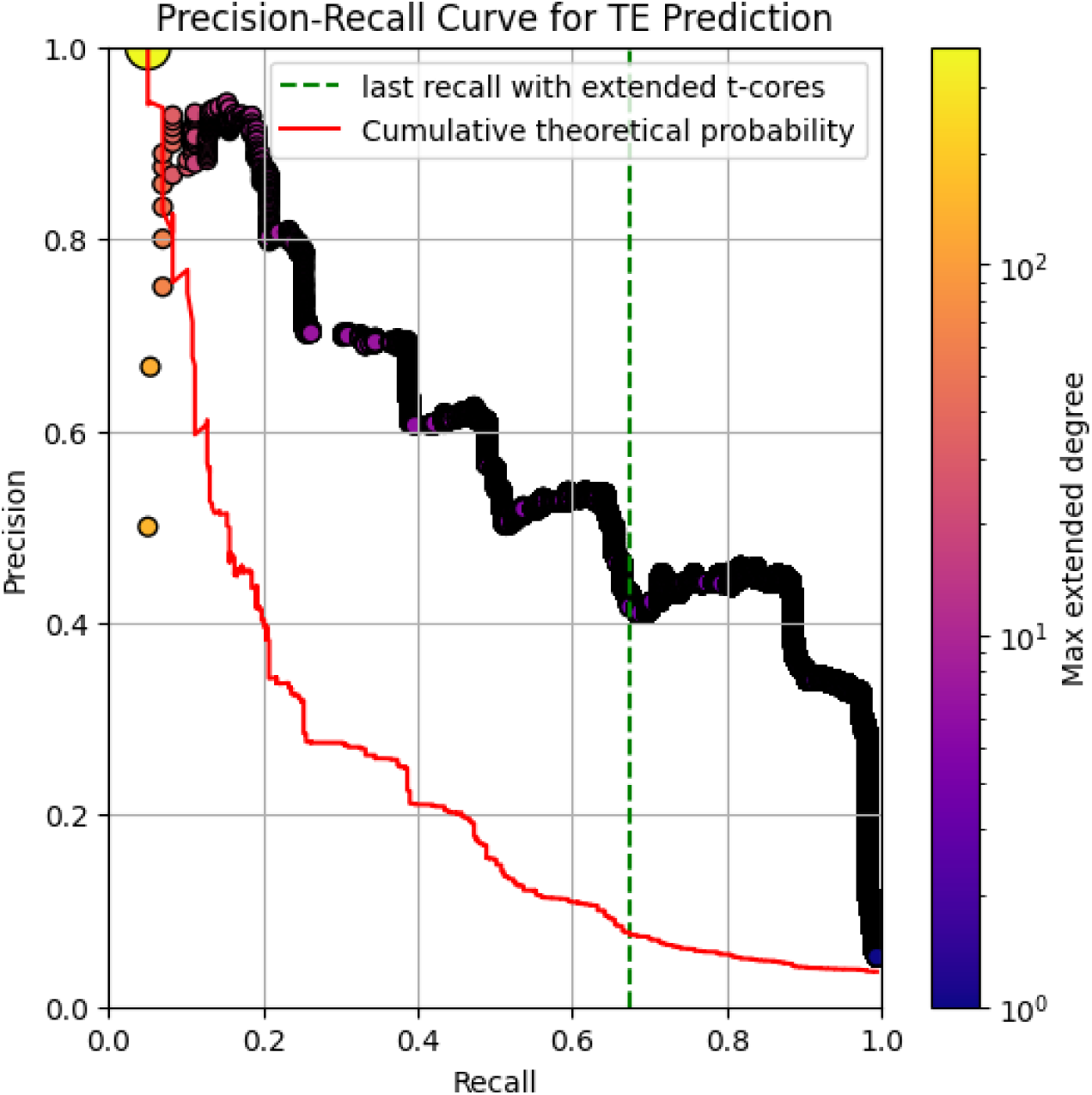
Cumulative performance of the extended *t*-core algorithm in SENSITIVE mode (*t* = 5) for the *Mus musculus* dataset. The graph shows the cumulative precision and sensitivity specifically for cores flagged as "Potential TEs", ordered by descending maximal extended degree. Dot sizes are proportional to the number of unitigs in each core. The red line represents the baseline performance expected from a random selection of unitigs, and the vertical green dashed line indicates the threshold *t* = 5.

As expected, the SENSITIVE mode allows for a much higher recall of expressed TEs compared to the PRECISE mode, although this gain in sensitivity comes with a moderate increase in false positives, particularly when considering all Potential TEs cores.

### *Canis lupus familiaris* extended *t*-cores

Running the extended *t*-core algorithm in PRECISE mode on the *Canis lupus familiaris* dataset yielded 364 distinct cores. As with the mouse dataset, it is crucial to emphasize that these results were achieved entirely *de novo*, without any reliance on a reference genome or prior knowledge of TE sequences.

Validating these results, however, presented a unique challenge: the Dfam database currently lacks a curated library of TEs specific to the dog, and RepBase consensus sequences are subject to licensing restrictions that prevent free redistribution. To overcome this limitation, we utilized the UCSC RepeatMasker annotation track for the dog reference genome assembly (canFam4) (Bao et al., 2015; Jurka, 1998, 2000; Jurka et al., 2005; Smit et al., 1996; Smit, 1996, 1999). For each annotated TE family, we randomly sampled 100 genomic copies directly from the reference genome. This strategy provided a robust substitute for formal consensus sequences, capturing the natural intra-family sequence diversity required to validate the identified *t*-cores, while avoiding seeding failures caused by an excessively high copy number of TEs in the genome (some TE families have more than 100,000 genomic copies). Moreover, cores initially flagged as false positives were manually inspected to determine their true biological origin.

Out of the 364 distinct extended *t*-cores, 243 were assigned to the "Potential TEs" category. Within this subset, only six cores were confirmed as false positives (originating either from a sequencing contamination or from a gene), leading to a precision of 97.5%. This result highlights the strength of our *de novo* approach: even though these cores could not be validated using standard Dfam consensus sequences due to database incompleteness, our cores successfully isolated true TEs.

Beyond identifying isolated repetitive regions, our tool also computes the induced subgraph of the cores. To do so, we compute all the paths connecting distinct cores, without entering into any other core. Since our extended *t*-cores are designed to capture borders of TEs, the paths between cores may correspond to a transcript of a TE. Specifically, if two cores are connected by a high number of paths, they are designated as strongly linked neighbors and reported in the summary output file to allow further investigations.

One such example is depicted in Figure 6, illustrating the connectivity between Core 4 (classified as a microsatellite) and Core 75 (classified as a "Potential TE"). Several unitigs from both cores are aligned to one of the paths connecting them, on Figure 6.

**Figure 6.**
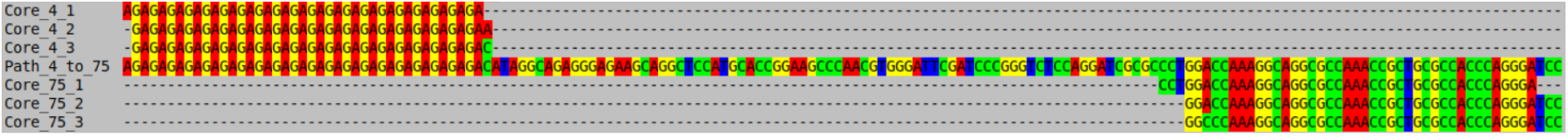
Potential reconstruction of a TE. Alignment of three unitigs (Core4_*i*) from the fifth extended *t*-core and three unitigs (Core75_*i*) from the 76th extended *t*-core with the path connecting the two cores (Path_4_to_75). Core 4 is flagged as the microsatellite (*AG*)^n^ whereas the core 75 is flagged as "Potential TE". The path Path_4_to_75 aligns to a *SINEC2A1_CF* TE

In this case, the path Path_4_to_75 aligns entirely to the *SINEC2A1_CF* TE, a SINE known to harbor an internal (*AG*)*^n^* microsatellite. This example demonstrates the power of combining extended *t*-cores with their induced subgraph to reconstruct and discover full-length TE transcripts *de novo*. Notably, approximately 31% of all connecting paths between extended *t*-cores in the cDBG are indeed part of a TE consensus. Most of the other connecting paths correspond to parts of transcribed genes containing two borders of different repeats. These repeats are connected because they co-occur in the same host gene.

### Benchmarking of the extended *t*-cores computation

Benchmarks were made using the *Mus musculus* dataset, which is composed of 52 million paired-end reads (145 bp long). Processing this entire dataset takes 24 minutes, generating a file footprint of approximately 22 GB. The complete workflow of our tool, *ET-core*, is depicted in Figure 7. By default, this pipeline includes a quality control and preprocessing step using *FastP* Chen et al., 2018, which accounts for 5 minutes of the total runtime.

**Figure 7.**
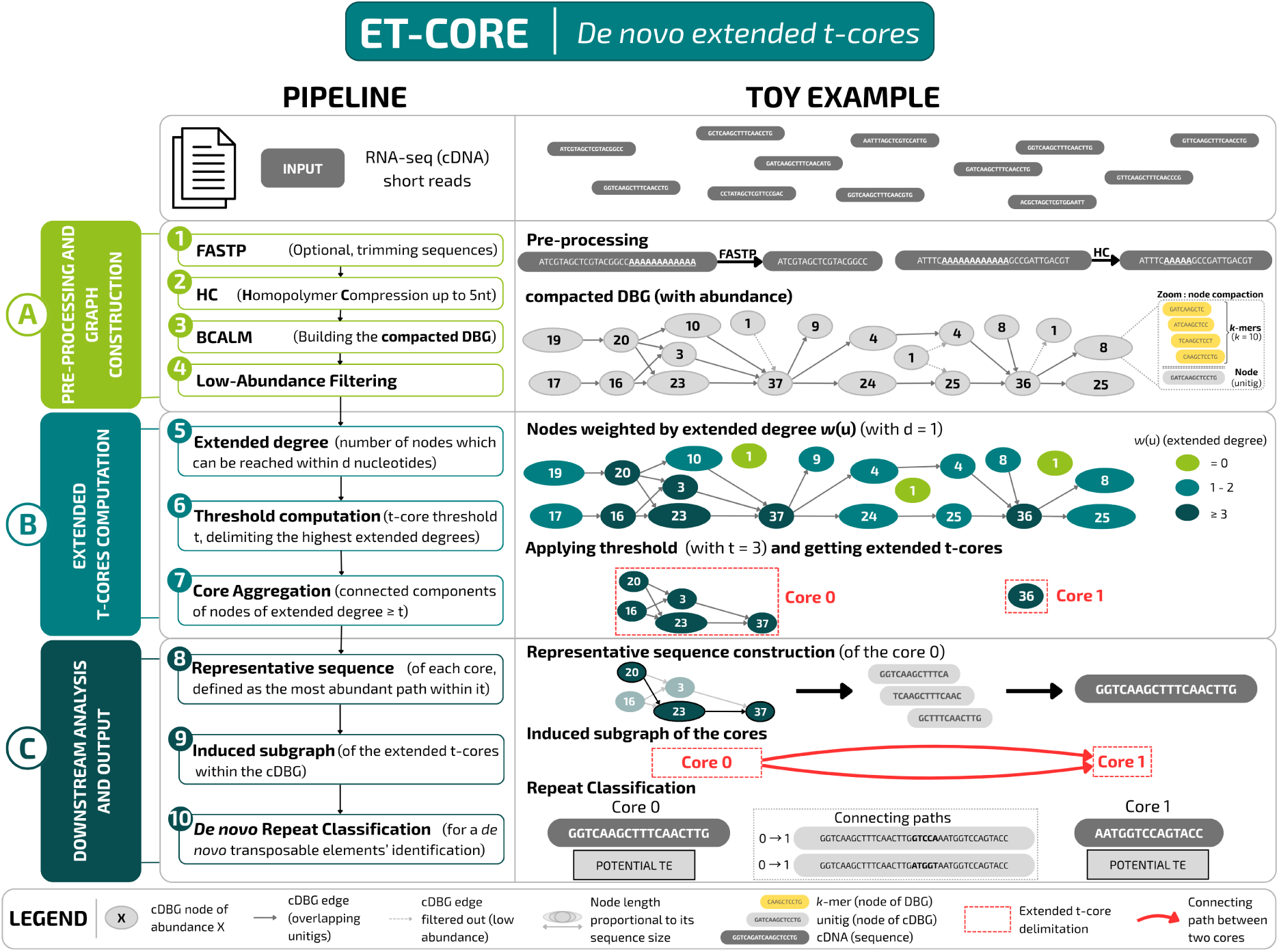
Overview of the *ET-core* pipeline. The flowchart illustrates the sequential steps of our tool, accompanied by a simplified toy example.

To evaluate the scalability of our approach, the preprocessed dataset was randomly down-sampled using *seqtk* (Li, 2012). These subsampled datasets were then processed by our tool with the preprocessing step disabled (–no-fastp). This was done to isolate the performance of the core graph algorithms and avoid any benchmarking bias introduced by the external *FastP* step. The resulting metrics for execution time, disk space, and memory usage are presented in Figure 8. Additionally, the number of unique TEs recovered from the extended *t*-cores at each sequencing depth is illustrated in the same figure.

**Figure 8.**
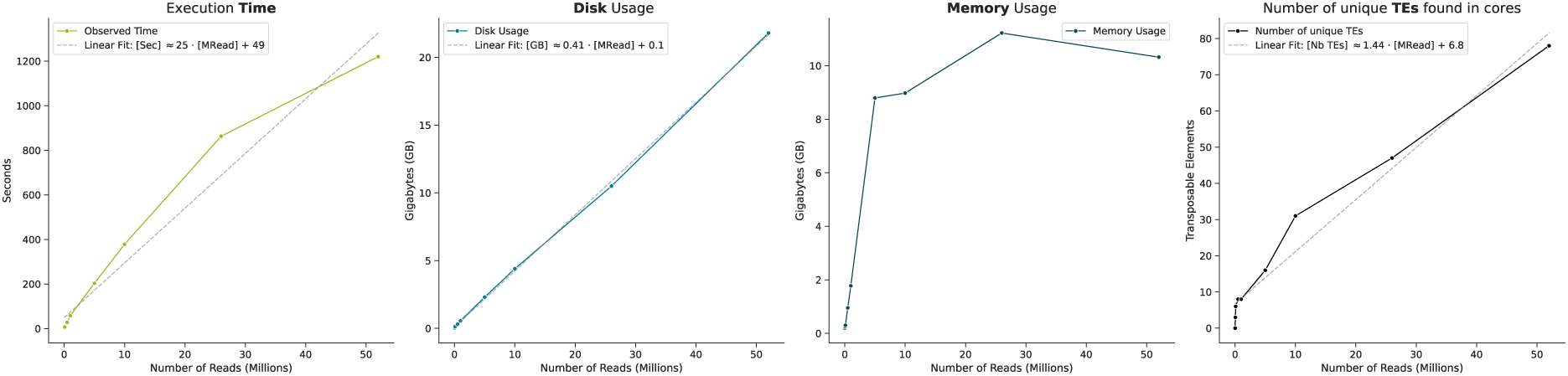
Benchmarks (Time, Disk Usage, Memory Usage and number of unique TEs found) of RNA-seq paired-ended samples (145bp long) from *Mus musculus* brain cells. Samples range from 10 thousand reads to 52 million reads.

Benchmark results confirm the linear scaling of both execution time and disk space usage, demonstrating that the tool is efficient enough to run on a standard laptop.

Regarding memory usage, the maximum footprint reaches a plateau as the number of reads increases. This is primarily because the construction of the cDBG by *BCALM2* is the most memory-demanding step of the pipeline, and its maximum memory usage can be strictly capped by a user-defined parameter (set to 14 GB by default). The subsequent steps required to compute the extended *t*-cores have a negligible memory footprint in comparison. Consequently, across all species analyzed in our diverse experiments, a limit of 14 GB of RAM was consistently sufficient to process and load the entire compact graph into memory.

Finally, the last curve, which depicts the number of unique TEs identified within the cores as a function of the number of reads, demonstrates the significant advantage of data sampling. Computing the cores using only one-tenth of the dataset takes approximately 3 minutes (compared to 24 minutes for the full dataset, see Table 1). Remarkably, this sampling strategy still enables the recovery of the 16 most highly expressed TEs without introducing a single false positive among the "potential TE" cores.

**Table 1.**
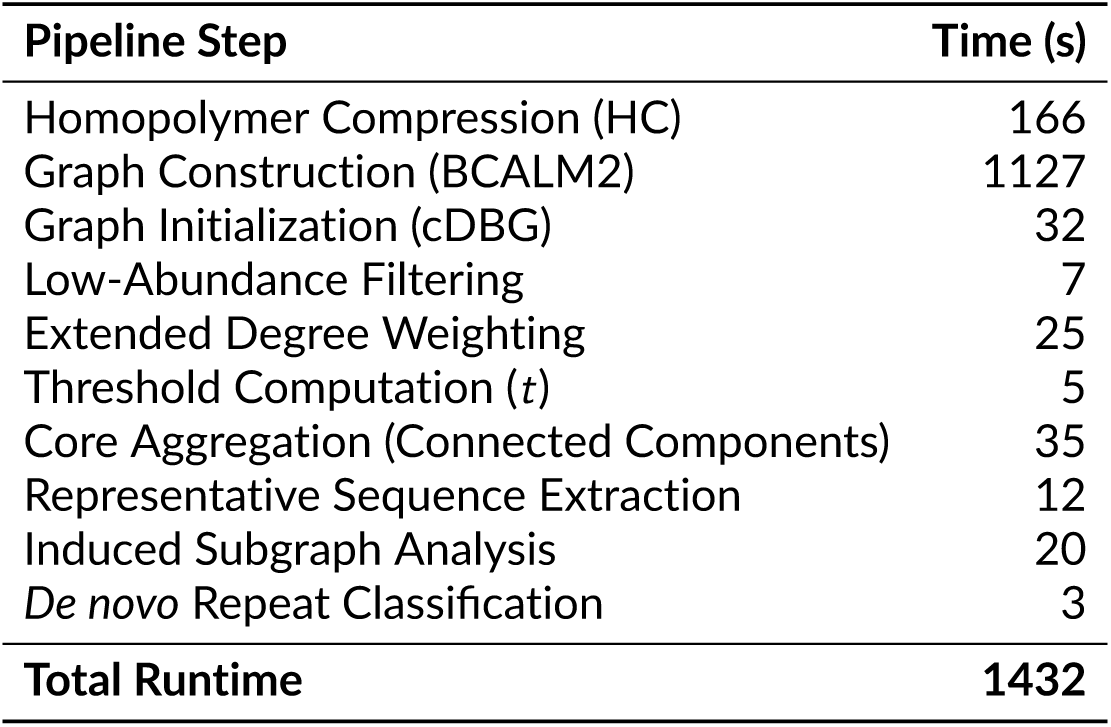
Execution time benchmark for each step of the extended *t*-core pipeline using the full dataset of *Mus musculus*.

| Pipeline Step | Time (s) |
| --- | --- |
| Homopolymer Compression (HC) | 166 |
| Graph Construction (BCALM2) | 1127 |
| Graph Initialization (cDBG) | 32 |
| Low-Abundance Filtering | 7 |
| Extended Degree Weighting | 25 |
| Threshold Computation ($t$) | 5 |
| Core Aggregation (Connected Components) | 35 |
| Representative Sequence Extraction | 12 |
| Induced Subgraph Analysis | 20 |
| *De novo* Repeat Classification | 3 |
| **Total Runtime** | **1432** |

## Discussion

In this paper, we introduce the notion of extended-t-core, which enables to capture very dense regions in graphs with bounded degrees. We apply this notion to DBG built from RNA-seq data in mouse and dog and show that cores can effectively be used as *de novo* predictors of TEs. Importantly, we do not use any prior information on the sequence of the TEs to identify them. We use only the structure of the DBG.

### The transcriptomic repeatome in a De Bruijn graph

Using extended *t*-cores as inexact repeats in the De Bruijn graph, we observed several remarkable features that could easily be overlooked with a classical De Bruijn graph approach.

Inexact repeats are predominantly expressed within host genes. Most inexact repeats identified in the transcriptome of mouse and dog are located within protein-coding genes and are co-transcribed with their host gene. While very few of these repeats are located within coding exons, many are within UTRs or introns. RNA-seq data should in principle not contain intronic sequences, but in practice, this is not the case and even when using stringent protocols which focus on poly(A)+ transcripts, there always remains a fraction of ∼ 5% of pre-mRNA (Tilgner et al., 2012). Our method can detect such intronic repeats, as long as they are present in very similar copies within genes that are sufficiently expressed. In practice, extended-t-cores often correspond to a mixture of exonic repeats and intronic repeats.

Intergenic repeats can also be detected. This is the case for instance of L1Md, which is known to be still active in mouse genomes. Active transposable elements are expected to be highly conserved. If only exact copies of a transposable element are present in the transcriptome, our method would be unable to detect them. The very fact that some copies that diverged are still transcribed, or co-transcribed together with their host gene, creates the possibility for our method to identify the structure. Out of all copies that are expressed, our method cannot decide which one, if any, is still active.

Of note, recent paralogs could in principle generate extended-t-cores. There are two reasons why this was rarely the case in our analysis. First, we used *k* = 41, which means that only very recent paralogs are concerned. Second, when computing the extended degree of a unitig, *k*-mers with a Hamming distance smaller than 2 are counted as one. While this refinement of the definition of extended degree was originally introduced to control for the impact of sequencing errors located in homopolymers, it also has the consequence of favouring the detection of repeats which differ by indels or their context of insertion. Autonomous repeats which only differ by a few substitutions will therefore not be reported.

Repeat clusters form large connected components. UTRs and introns can accumulate multiple distinct inexact repeats with minimal functional impact or selective pressure. Consequently, these repeats are often located in close proximity to one another along the same transcript, leading to their entanglement within the De Bruijn graph. This biological clustering results in the formation of large giant components in the graph. For instance, in our *Mus musculus* analysis, 60% of the identified extended *t*-cores were found to reside within a single, large connected component.

### Evolutionary interpretation of identified TEs

The most prominent elements in the extended *t*-cores of the *Mus musculus* dataset consist of either transcriptionally active TEs (e.g., L1Md, found in the core 40, see Figure 9) or relatively young TEs with an exceptionally high genomic copy number (e.g., SINE B superfamily) (Kawase and Ichiyanagi, 2023). In all cases, the top-ranked TEs identified by our method belong to families characterised by a high copy number, for which remnants of TEs may inflate the transcription signals.

**Figure 9.**
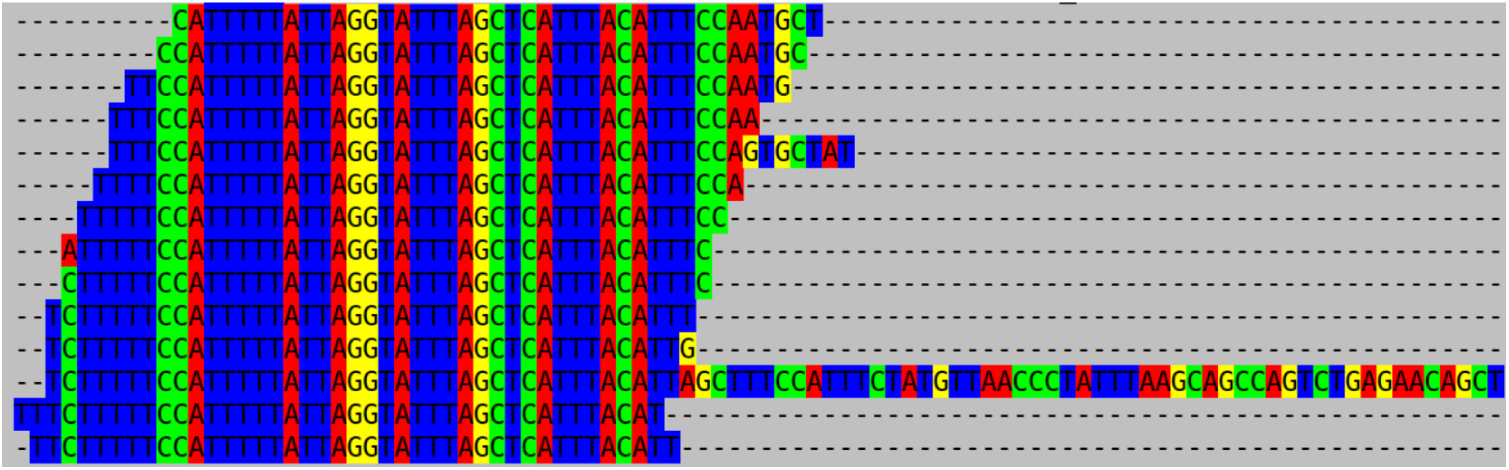
Alignment of some unitigs of an L1Md core.

Conversely, some TEs are expressed but remain undetected by our method, even using the SENSITIVE mode. The majority of these false negatives consist of low-copy ERV elements, being transcribed outside genes.

A notable example in the *Mus musculus* dataset is the *RLTR4_MM* / *RLTR4_MM-int* element. It is a young element (1%–2% divergence) transcribed with a coverage depth of 400 and which is not captured in an extended *t*-core. This omission is a consequence of two factors. First, the unitigs associated with *RLTR4_MM* have a maximal extended degree of 4, less than the threshold *t* = 5 used for the SENSITIVE mode, causing them to be filtered out during the *t*-core extraction. Second, this element is still transcriptionally active and is mainly transcribed outside genes, preventing its unitigs from having high extended degrees according to the definitions (see section Methods).

Altogether, these results demonstrate that our approach is highly effective at recovering young, not necessarily active, TEs with high copy numbers, making it a valuable tool for exploring the repeatome of non-model species.

### SINEs and LINEs in mammals

In mammalian genomes, Short Interspersed Nuclear Elements (SINEs) and Long Interspersed Nuclear Elements (LINEs) represent the two most prominent families of TEs. The retrotransposition of these elements is deeply intertwined: SINEs (such as *Alu* in humans or *SINEC* in canines) are non-autonomous and systematically hijack the enzymatic machinery encoded by active LINEs (specifically LINE-1) to mobilize and replicate throughout the host genome (Dewannieux et al., 2003; Halo et al., 2021; Kajikawa and Okada, 2002).

This dynamic is particularly well characterised in *Mus musculus*, which harbors two major active TE lineages: the L1Md family (e.g. the core 40, see Figure 9, Zhou and Smith, 2019), and the SINE B2_Mm1a subfamily Ichiyanagi et al., 2021). Notably, experimental evidence has demonstrated that the transcription and retrotransposition of *B2_Mm1a* subfamilies are strictly dependent on the reverse transcriptase provided by L1Md elements (Dewannieux et al., 2003; Dewannieux and Heidmann, 2005; Kajikawa and Okada, 2002; Ohshima and Okada, 2005).

Remarkably, standard genomic mapping strategies completely fail to identify *SINEC* elements in the dog transcriptome. Their extremely high copy number in the genome causes mappers to report them as unmapped, even using the most sensitive options. Indeed, mappers usually use a seed to find a small number of possible gene loci and then try all possible alignments at those loci. Yet, *SINEC* are so abundant that all candidate seeds match too many loci, making mappers blind to such elements.

### Proposition of substructures in TE families

Using the PRECISE mode of our tool, we observe that the extended *t*-cores are dominated by SINEs and LINEs. *B2* elements are found in approximately 20% of all *Mus musculus* cores, while *SINEC* elements are associated with roughly 80% of the *Canis lupus familiaris* cores.

More precisely, we can actually report several extended *t*-cores for a single TE consensus family, as depicted in Figure 3. In this example, the core 0 perfectly matches the *B2_Mm1a* element, and the cores 10 and 12 harbor one different mutation over the same site: a CG site is mutated into a CA (in core 10) and into a TG (in core 12).

These specific point mutations could be evidence of a biological transition in the evolution of the SINE element. As pointed out by Ichiyanagi et al., 2021, *B2_Mm1a* copies (represented by Core 0) are typically CpG-rich. The host genome utilises these CpG dinucleotides as targets for DNA methylation, silencing the element’s transcription. CpG sites are hotspots of base substitutions during evolution, and when the total CpG count drops to 2 or fewer, the recruitment of the silencing is altered, resulting in the loss of epigenetic control (Ichiyanagi et al., 2021).

The identification of these distinct substructures by our tool stratifies the ***B2_Mm1a*** family into biologically different subpopulations

Core 0 captures an intact, active element. Conversely, Cores 10 and 12 capture two alternative copies that diverged from the consensus copy.

Another example of a TE copy which lost several of its CpG sites with respect to the consensus sequence is B2_Mm2 which corresponds to core 1. This might explain its transcriptional success, producing more than half of all B2 SINE RNA in spermatogonia (Ichiyanagi et al., 2021).

### Why not use long reads?

While third-generation long read sequencing technologies (such as PacBio and Oxford Nanopore) are increasingly becoming the standard for genome assembly, our method deliberately focuses on short-read RNA-seq data for two fundamental reasons.

First, short-read sequencing remains the most cost-effective and widely adopted technology. Over the past decade, the scientific community has generated millions of short-read RNA-seq datasets across diverse conditions, tissues, and species, which are now freely available in public databases like SRA. Our tool is designed to unlock this massive, untapped historical data, allowing researchers to explore the repeatome without the need for new sequencing runs.

Second, the accurate detection of repeats requires both low sequence error rates to distinguish between distinct copies and high coverage depth. However, long read sequencing typically yields significantly shallower coverage than short-read sequencing. To illustrate this, we analysed long read sequences from the same *Mus musculus* individual (one ONT RNA dataset, SRA accession ERR2680375 and one ONT cDNA dataset, SRA accession ERR2680377). We aligned the reads on the TE sequence consensus using Minimap2 (Li, 2018). With the ONT RNA dataset, we recovered only 33 consensus elements, ten times fewer than those identified using short-read data. Using the cDNA dataset, we only recovered 163 elements, representing less than half of the short-read results. Consequently, due to its unmatched coverage depth, short-read RNA-seq remains the most robust strategy for comprehensively capturing the high diversity of TE expression levels. Even when the consensus are known, this mapping approach is not as powerful simply because the depth is less.

## Further Work and Applications

Looking forward, the extended *t*-core framework opens several promising avenues for future research. While our current study focused on a single diploid individual, it could be extended to pooled RNA-seq datasets. Specific developments would however certainly be required to model gene or TE polymorphisms at the population level. Furthermore, the framework could be extended to the analysis of complex polyploid genomes. Finally, a key future development will involve leveraging the induced subgraph of the *t*-cores to improve the *de novo* identification of TEs and to automatically assemble full-length TEs.

## Supporting information

Extended t-cores summary of the Mus musculus data

## Acknowledgements

The authors would like to thank Thomas Derrien and Rita Rebollo for their advice and insightful discussions.

## Data, script, code, and supplementary information availability

Datasets are available online (https://www.ncbi.nlm.nih.gov/sra[SRA NCBI]) under the accession numbers SRR15254973 (*Canis lupus familiaris* RNA-seq), ERR2680378 (*Mus musculus* RNA-seq), ERR2680375 (*Mus musculus*, ONT RNA), ERR2680377 (*Mus musculus*, ONT cDNA); Tools, scripts, code and reproducibility are available online (https://github.com/sdarmon/ET-core[ET-core]);

A released version of the tool is available under the following DOI url https://doi.org/10.5281/zenodo.21219806;

